# Potentials of Aqueous Extract of *Bambusa vulgaris* to Reactivate β-Cells of Pancreas in Alloxan-Induced Diabetic Wistar Rats

**DOI:** 10.1101/2020.04.11.037499

**Authors:** MO. Akiibinu, OS. Folawuyo, IO. Oyekan, I. Ibikunle, OT. Kolawole, A. Adesiyan, J. Anetor, SO. Akiibinu, F. Tayo

**Affiliations:** Department of Chemistry and Biochemistry, Caleb University Lagos, Nigeria; Department of Medical Laboratory Sciences, Ladoke Akintola University of Technology, Ogbomoso, Oyo State, Nigeria; Department of Pharmacology, Ladoke Akintola University of Technology, Ogbomoso, Oyo State, Nigeria; Department of Chemical Pathology, University of Ibadan, Ibadan, Nigeria; Pharmaco-Herbal Development Laboratory, Famot HealthCare Ltd, Iyeri Estate, Ikorodu-Itoikin Road, Imota, Lagos State; Nursing Services Department, University College Hospital Ibadan, Nigeria

**Keywords:** β-cells reactivation, glucose metabolism, *Bambusa vulgaris*

## Abstract

Ability of the damaged pancreas to generate new β-cells when activated with external stimuli has been documented. This study assessed the potentials of aqueous extract of *bambusa vulgaris* leaf to reactivate damaged β-cells in insulin dependent diabetes mellitus (IDDM). Eighteen healthy male Wistar rats (12 weeks old; weight= between 150 and 200g) were used for this study. The rats were randomly divided into three groups (six per group); group A (un-induced); group B (untreated alloxan-induced diabetics); group C (alloxan-induced diabetics treated with 200mg/kg body weight of freshly prepared extract of *bambusa vulgaris* leaf). Fasting blood sugar (FBS) and plasma insulin levels were determined in all animals using glucometer and enzyme linked immunosorbent assay methods respectively. Sections of pancreas tissues were prepared for histology. IDDM was confirmed in groups B and C (FBS increased significantly=p<0.05) after 2 days of alloxan administration). The FBS remained significantly (p>0.05) higher in group B, compared to group A, but reduced significantly (p<0.05) in group C after 7 days of treatment with *bambusa vulgaris* leaf extract. On the 7th day, plasma insulin level decreased significantly (p<0.05) in group B, but no significant difference observed in group C compared with group A. Histology reports showed damaged pancreas in group B, while Group C showed normal islet cells after 7 days of oral administration of *bambusa vulgaris* extract. In conclusion, aqueous extract of *Bambusa vulgaris* could restore the activities of alloxan-damaged pancreas. The extract could be a reliable alternative to synthetic pharmaceuticals in the treatment IDDM.

## Introduction

Type 1 diabetes mellitus is characterized by autoimmune inflammatory process which selectively affects the pancreatic islets causing the destruction of β-cells of pancreas by macrophages, CD4+ and CD8+ T cells infiltrating the islets [1, 2]. The β-cells can be selectively destroyed by an inflammatory process characterized by presence of immuno-competent and accessory cells in the infiltrated pancreatic islets. Tumor and viral (Coxsackie B virus) factors have also been implicated in the pathogenesis of IDDM [3]. The dysfunctional β-cell of the pancreas is the basis of the insulin therapy to maintain normoglycaemia in IDDM [4]. Studies show that the dysfunctional β-cell can be reactivated by increasing the mass of existing β-cells in the pancreas [5], when extra-islet cells differentiate into β-cells in response to external factors [5, 6, 7]. Another reports show that regulation of plasma glucose could still be achieved by the regeneration, proliferation, and replication of pre-existing β-cells, as well as by the neogenesis of β-cells from precursor or stem cells or by trans-differentiation from other differentiated cells [8, 9]. Many hypotheses support possibility of trans-differentiation of β-cells [10]. In animal models, insulin dependent diabetes mellitus can be induced by administration of streptozotocin or alloxan. Streptozotocin is a glucose analogue with methylnitrosourea that can modify biological macromolecules, fragments DNA, impairment of the signalling function of β-cell mitochondrial metabolism and destroys the β-cells of the pancreas [11]. Alloxan is a toxic glucose analogue that preferentially accumulates in pancreatic β-cells via the GLUT2 glucose transporter. In the presence of glutathione and other intracellular thiols, alloxan undergoes cyclic redox reaction to generate reactive oxygen species (ROS) and dialuric acid [12]. Auto-oxidation of dialuric acid generates superoxide radicals, hydrogen peroxide and hydroxyl radicals with fatty degeneration, signs of mitochondrial damage, perivascular edema and leukocytic infiltration [13]. The hydroxyl radicals generated ultimately destroy the beta cells, cause a significant deficit in pancreatic β-cell mass, selectively inhibit glucose-induced insulin secretion, leading to insulin dependent diabetes mellitus [14].

*Bambusa vulgaris*, known as Bamboo (English) is the fastest growing perennial evergreen arborescent plant belonging to the true grass family Poaceae, subfamily Bambusoidae and the branches can be as long as 11m (35ft) [15]. The plant grows naturally in China, India [16] and other regions like Africa, Latin America, and various oceanic islands [17]. In Nigeria, *bambusa vulgaris* is called Oparun amongst the Yorubas, Atosi amongst the Ibos, Gora amongst the Hausa and Iko amongst the Bini. The human consumption of *Bambusa vulgaris* dates back to antiquity. It is used for shelter, fuel, transportation, food, musical instruments, arts, craft, and medicines [18]. The leaves of the genus *Bambusa* has been shown to possess diverse therapeutic properties such as antioxidant, antimicrobial (antibacterial, antifungal, antiplasmodial, etc.), antidiabetic and anticancer effects [19]. The leaf decoction is consumed orally to treat diabetes traditionally by the Moran folk of Tinsukia district of Assam, India [20]. In vitro antidiabetic studies of the leaves *Bambusa vulgaris* have shown promising results [21]. Earlier reports show that the aqueous leaf extract is rich in flavonoids (0.05%), alkanoids (4.10%), tannins (0.93%), saponins (1.14%), phenolics (2.27%), glycosides (0.63%) and anthraquinones (0.06) [22]. Results of the toxicity test carried out by Senthilkumar *et al*. [23] on the *Bambusa vulgaris* showed no sign of toxicity up to the dose of 4000mg/kg body weight. Since insulin replacement is the only treatment strategy in the management of IDDM, investigating the possibility of aqueous extract of *Bambusa vulgaris* to reactivate the damaged β-cells of the pancreas in alloxan-induced diabetic rats provided a basis for this study. This study was designed to assess the ability of aqueous extract of *Bambusa vulgaris* to reactivate the β-cells of pancreas in alloxan-induced diabetic Wistar rats.

## MATERIALS AND METHODS

### Materials

#### Plant material collection and authentication

Fresh and matured bamboo leaf was harvested from Osogbo Forest Reserve, Isale-Osun, Osogbo, Osun State, Nigeria. The specimen was identified and authenticated at the herbarium of the Faculty of Pharmacy, Obafemi Awolowo University, Ile-ife, Osun State, Nigeria and a voucher specimen number, FA2254 was deposited.

#### Preparation of Plant Extract

A 500g of air-dried leaves of *Bambusa vulgaris* was rinsed twice in 1000 ml of distilled water to remove dust and adhering dirt. This was then transferred into 2000 ml of hot distilled water maintained at 80^0^C for 45 minutes to remove some unwanted adherents and the leaves rinsed with another 1000 ml of hot (80°C.) distilled water. The extract was pulverized with glass pestle and mortar before passing through a 250 µl mesh sieve, and then freeze-dried. The powdered extract was stored in a chamber of activated descicator for 48 hrs and then transferred into air-tight, amber colored screw capped bottle and stored at -20^0^C until needed [24].

#### Experimental Animals

Eighten (18) male healthy Wistar rats (12 weeks old) obtained from the Experimental Animal Unit of the Department of Medical Laboratory Science, Ladoke Akintola University of Technology, Mercyland Campus, Osogbo were used for this study. The rats weighed between 150 and 200grams and maintained in galvanized wire mesh cages, freely ventilated and naturally illuminated animal rooms under hygienic conditions. They were made to acclimatize to the animal house condition for two weeks under laboratory conditions maintained at a temperature of 25°C and humidity of 50%. The animals were maintained on standard commercial pelleted rat feed and also provided with clean water. The cages were cleaned daily and washed weekly. All rats received humane care as stated in the “Guide for the care and use of laboratory animals” [25].

#### Preparation of Alloxan Solution

**1.5g of** alloxan monohydrate that has not been exposed to light was dissolved in 10ml of 0.9% NaCl solution (physiological saline). The salt was carefully weighed under minimum illumination to prevent its conversion to a toxic product.

#### Experimental Induction of Diabetes Mellitus in Wistar Rats

After an overnight fasting, blood samples were collected from the tail veins of all the Wistar rats for the determination of basal levels of blood glucose (FBS) before a single dose of 150 mg/kg body weight of alloxan monohydrate solution was injected intra-peritoneally into Wistar rats in groups B and C as described by Nwaka *et al.* [12]. Since alloxan is capable of producing fatal hypoglycaemia as a result of a transient massive pancreatic insulin release in response to alloxan attack, the rats were treated with 2ml of 10% glucose solution using orogastric tube 4 hours after induction (12). Diabetes mellitus was confirmed in groups B and C on 2nd day as reported by Ezekwe *et al.*, (26). Blood samples were also collected from the tail vein after 3rd, 4th, 5th and 7th day of alloxan administration for the determination of FBS levels.

#### Oral Administration of Aqueous Extract of *Bambusa Vulgaris*

Only the Wistar rats in Group C received continuous daily oral dose of 200 mg/kg body weight of freshly prepared aqueous extract of *Bambusa vulgaris* for a period of seven days. After the 7th day, the Wistar rats were bled using cardiac puncture technique and the blood sample put in ethylene diamine tetra acetic acid bottle for the determination of plasma insulin level. All Wistar rats were sacrificed and the pancreas removed for histology tests.

#### Determination of Plasma Insulin and Blood Glucose Levels

Plasma level of insulin was determined by using a commercially prepared kit (CUSABIO Rat insulin ELISA kit, Catalog number-CSBE05070r). Fasting blood glucose was determined by using a digital glucometer (Fine Test Auto-coding Premium, OSANG Healthcare Co., Ltd, Korea).

#### Histopathological Studies

The pancreas of the sacrificed Wistar rats from groups A, B and C were subjected to postmortem gross examinations. The pancreatic tissue removed from each Wistar rat was washed in iced cold saline immediately. A portion of the tissue was fixed in 10% neutral formal saline solution for histological studies. After fixation, the tissue was embedded in paraffin, solid sections were cut at 5 mm and stained using Heamatoxylin and Eosin stain as described by Strate et al. [27]. The slides were viewed at magnification of X 250 and photomicrographs taken. The pathologist in the Ladoke Akintola University Teaching Hospital interpreted the results.

### Statistical Analysis

Statistical analysis was carried out using Statistical Package for Social Sciences (SPSS) version 20 to compare data obtained from all groups using Student’s T test and ANOVA. Data were presented as mean ± standard deviation (STD). P-values<0.05 were considered statistical significant.

## RESULTS

As shown in Figure 1, there were significant (p<0.05) changes in the plasma levels of insulin when alloxan-induced diabetic Wistar rats (group C) were treated for 7 days with 200 mg/kg body weight of extract of *Bambusa vulgaris* compared to the untreated alloxan-induced diabetic Wistar rats (group B). There was no significant (p>0.05) difference in the level of plasma insulin in group C compared to group A (controls). As shown in Table 1 and Figure 2, the level of fasting blood glucose increased significantly in group B compared to group C. Meanwhile, the FBS level initially increased in group C and decline significantly (p<0.05) after 7 days of treatment with 200 mg/kg body weight of extract of *Bambusa vulgaris.* Photomicrograph of section (Figure 3) from pancreas tissue of untreated alloxan induced diabetic rats (group B) shows significantly destroyed exocrine glands and islet cells areas with some necrotic areas and biochemical changes affecting the staining outcome compared to group A (control) and group C. Meanwhile, the diabetic rats treated for 7 days with 200mg/kg body weight of aqueous leaf extract of *bambusa vulgaris* (groups C) show normal islet cells with mildly dilated blood vessels when compared with control.

**Table 1:**
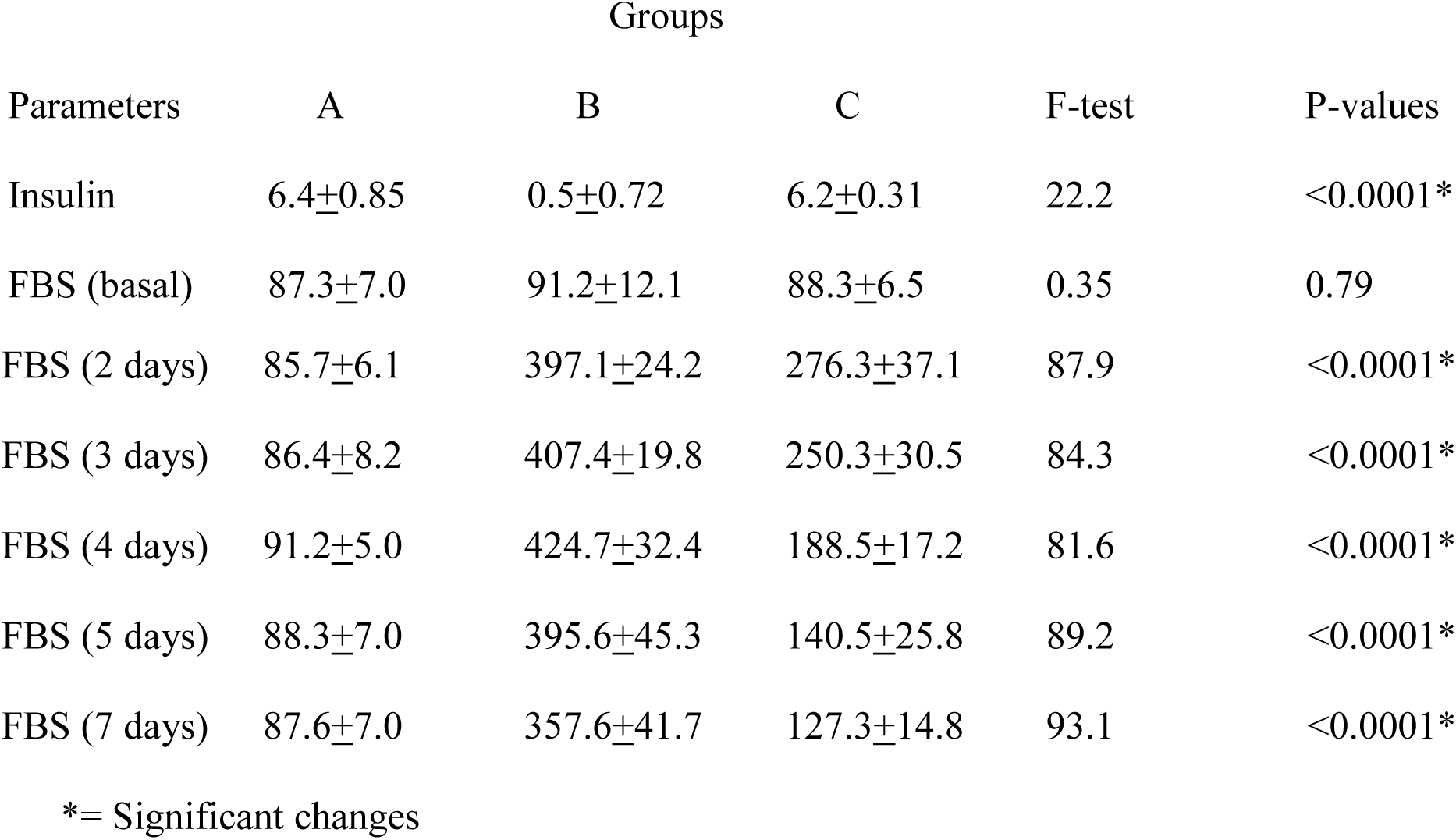
Comparison of Levels of Insulin (ng/ml) and Fasting Blood Sugar (mg/dl) in all Groups Using ANOVA

| Parameters | Groups |  |  | F-test | P-values |
| --- | --- | --- | --- | --- | --- |
|  | A | B | C |  |  |
| Insulin | 6.4±0.85 | 0.5±0.72 | 6.2±0.31 | 22.2 | <0.0001* |
| FBS (basal) | 87.3±7.0 | 91.2±12.1 | 88.3±6.5 | 0.35 | 0.79 |
| FBS (2 days) | 85.7±6.1 | 397.1±24.2 | 276.3±37.1 | 87.9 | <0.0001* |
| FBS (3 days) | 86.4±8.2 | 407.4±19.8 | 250.3±30.5 | 84.3 | <0.0001* |
| FBS (4 days) | 91.2±5.0 | 424.7±32.4 | 188.5±17.2 | 81.6 | <0.0001* |
| FBS (5 days) | 88.3±7.0 | 395.6±45.3 | 140.5±25.8 | 89.2 | <0.0001* |
| FBS (7 days) | 87.6±7.0 | 357.6±41.7 | 127.3±14.8 | 93.1 | <0.0001* |

\*= Significant changes

**Figure 1:**
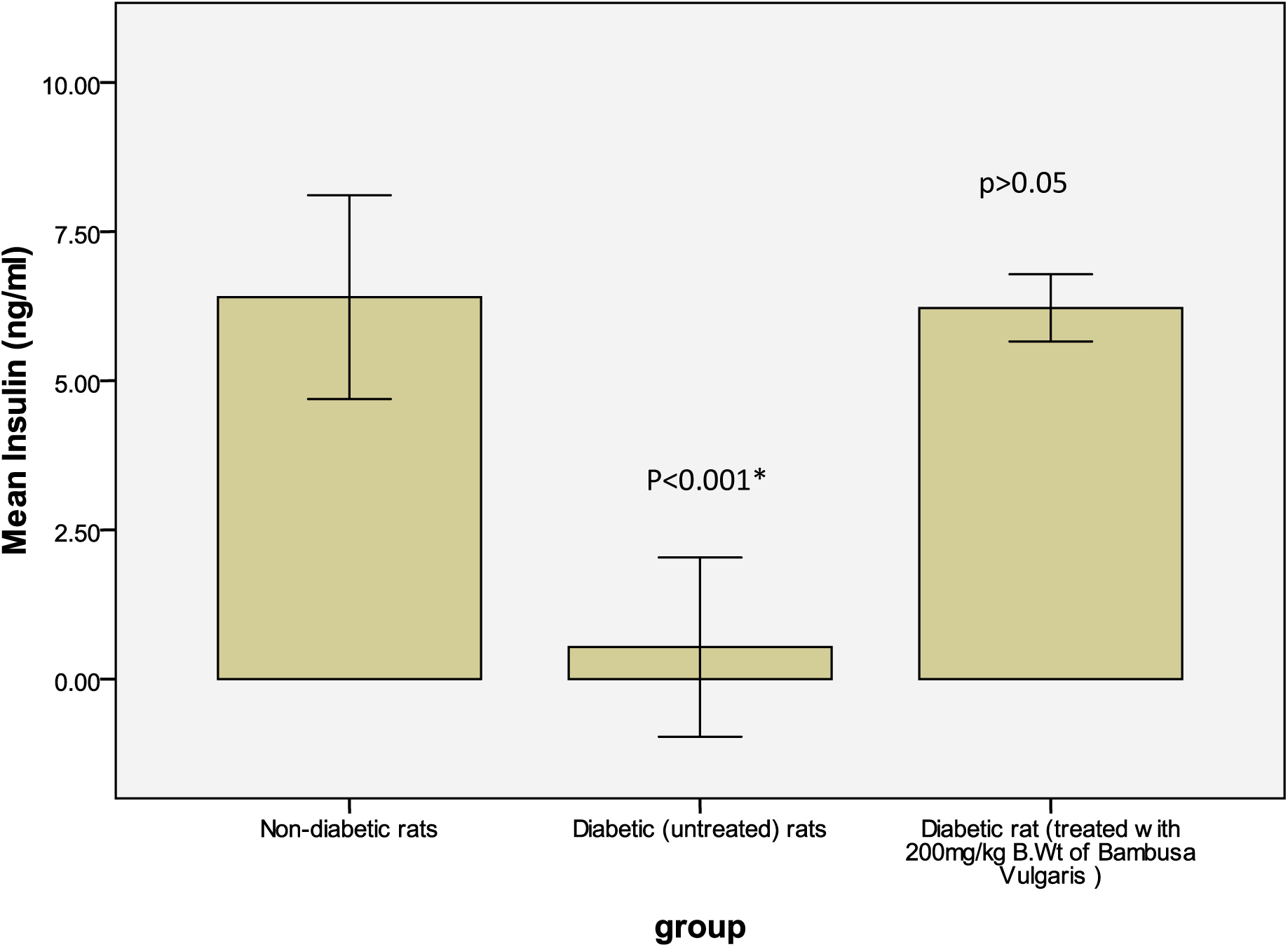
Plasma levels of Insulin in the Treated and Untreated Diabetic Wistar Rats on Day 7 of Experiment *= Significantly different from controls.

**Figure 2:**
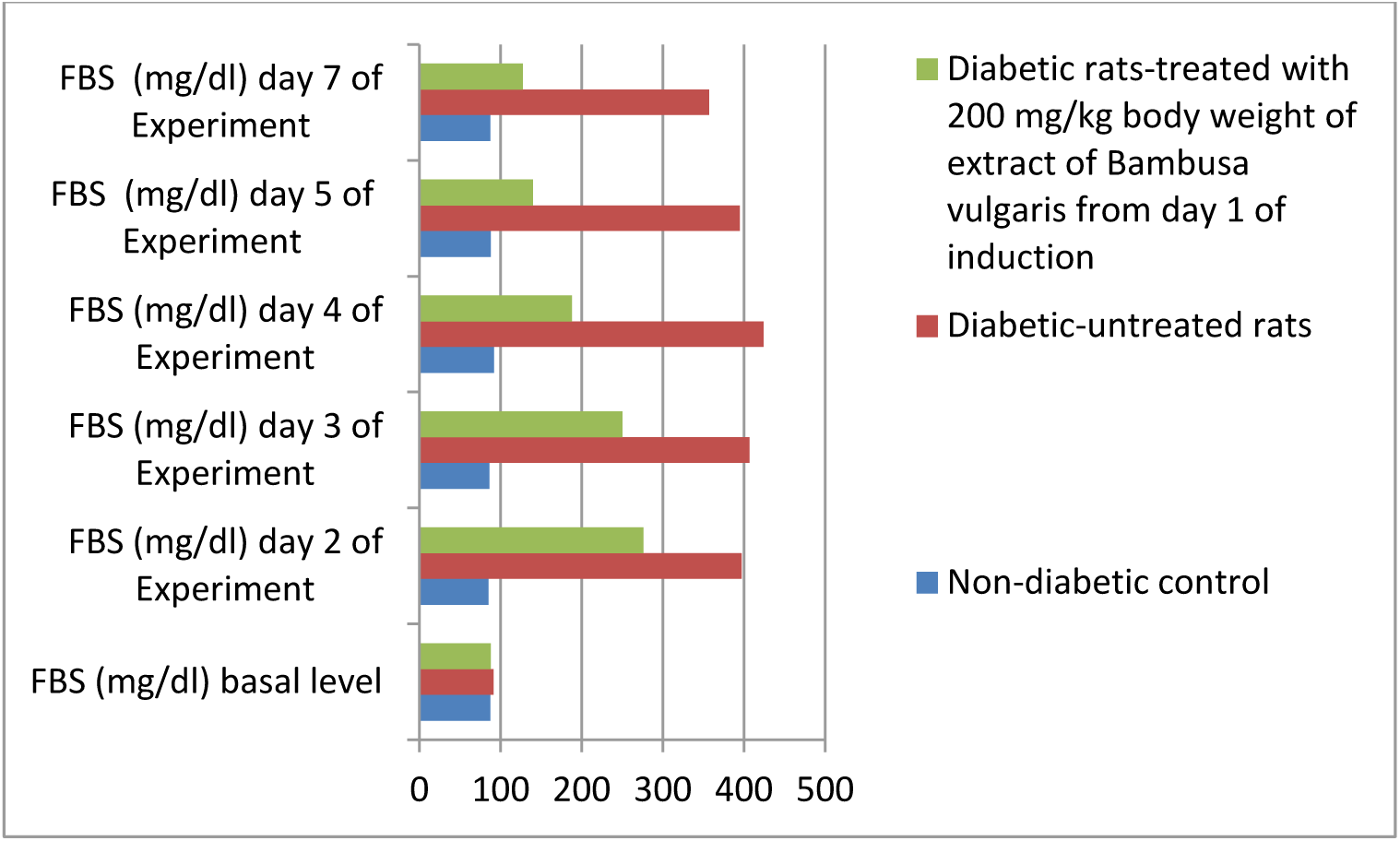
Changes in the Mean Levels of Fasting Blood Sugar in the Controls, Treated and Untreated Diabetic Wistar Rats.

**Figure 3:**
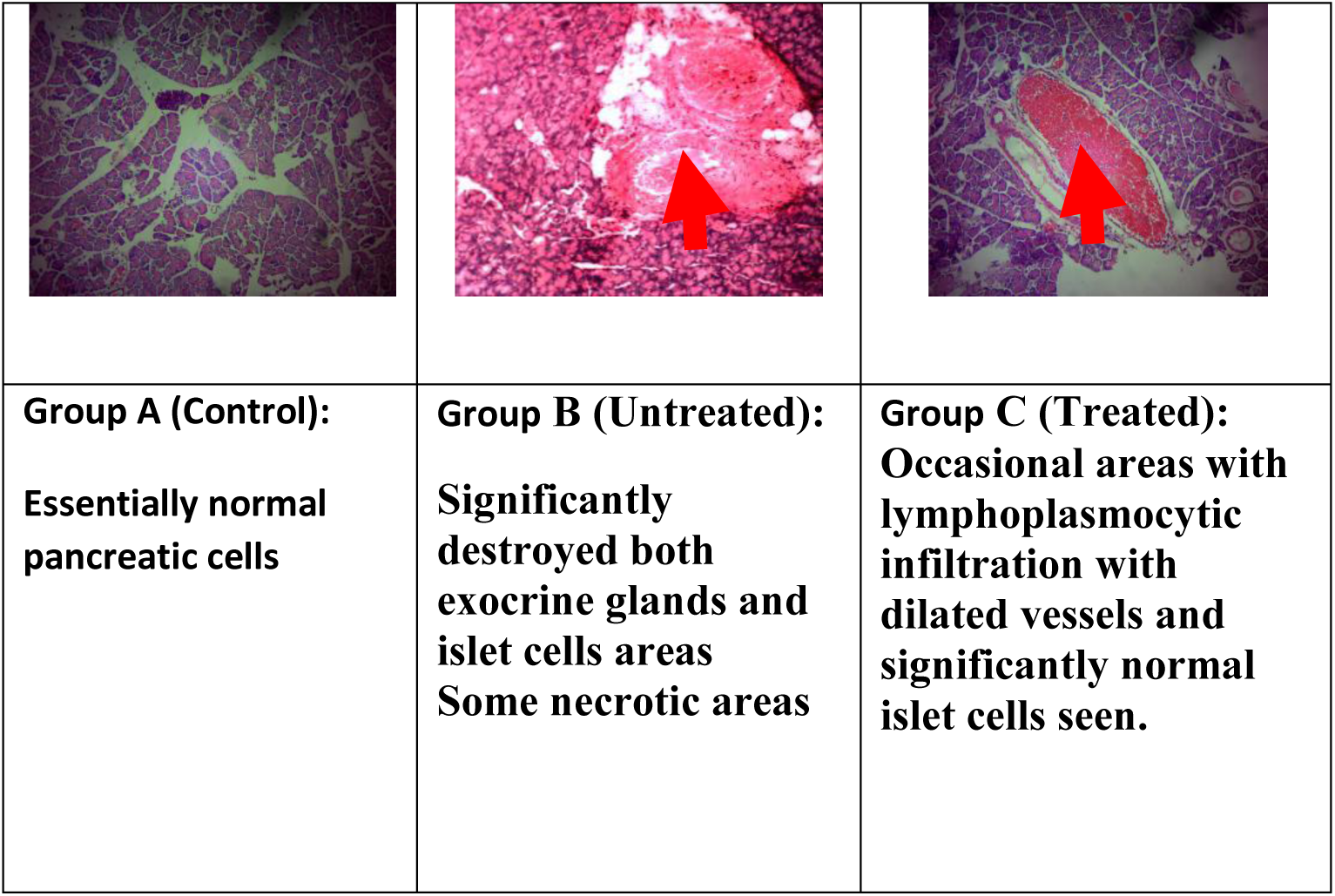
Histology of Pancreas of Treated and Untreated Alloxan Induced Wistar Rats.

## Discussion

Insulin dependent type is the most deadly form of diabetes mellitus. Destruction of β-cells by either autoimmune process or infection usually leads to absolute insulin deficiency [4, 14]. The β-cell insufficiency induced in animal models by using alloxan or streptozotocin is characterized by endothelial dystrophy, desquamation, transendothelial migration of immune and inflammatory cells into the pericapillary space, and the endocrine tissues and production of free radicals, leading to progressive injury of the β-cells [11]. The present study also shows the destructive effects of the administered alloxan on the pancreas of the Wistar rats. This agrees with the report of Latsabidze et al [6] who observed atrophied, hypertrophied and hyperemic islet with disorganized, necrobiotic changes and reduction of β-cells in alloxan treated Wistar rats. Latsabidze et al. [6] reported that most of the extra-islet cells including exocrine and endocrine granules specific for insulin secreting β-cells can potentially differentiated into β-cells when induced by external stimuli. They demonstrated that Plaferon LB promotes maturation and differentiation of extra-islet cells and takes part in the renewal processes of β-insulocytes in the pancreas of Alloxan/Plaferon LB-treated group. In the present study, the pancreases of those untreated alloxan-induced diabetic Wistar rats were significantly destroyed. Meanwhile, the alloxan induced diabetic Wistar rats treated with 200mg/Kg body weight of extract of *Bambusa vulgaris* leaf showed relatively normal pancreas, apart from occasional areas with lymphoplasmocytic infiltration and dilated vessels noticed. This study therefore hypothesize that aqueous extract of *Bambusa vulgarishas* may have potentials to reactivate damaged pancreas in alloxan induced diabetic rats

This study is the first to demonstrate increased insulin level in alloxan induced diabetic Wistar rats treated with aqueous extract of *Bambusa vulgaris.* Our study showed that plasma insulin level waned off in the untreated allloxsn-induced diabetic Wistar rats; but the level of insulin remained normal in alloxsn induced diabetic Wistar rats treated with 200mg/Kg body weight of extract of *Bambusa vulgaris* leaf. The antioxidant property of the extract of *Bambusa vulgaris* could prevent oxidative damage of the pancreas in these treated diabetic Wistar rats. Our finding corroborates the report of Weir et al [5] that the dysfunctional β-cell can be reactivated by increasing the mass of existing β-cells in the pancreas. This study also agrees with a previous finding by Singh and Gupta [28] who observed regeneration of β-cells in the pancreas of alloxan diabetic rats treated with acetone extract of *Momordica charantia.* In another study on reactivation potential of pancreas cells by external stimuli, Latsabidze et al [6] reported that Plaferon LB can protect the insulin producing β-cells, reduce intensity of apoptosis, restore potentials of mitochondria and stimulate β-cell proliferation by increasing its mitogenic activity. Progressive reactivation of the pancreas in the Wistar rats treated with 200mg/Kg body weight of extract of *bambusa vulgaris* leaf and restoration of insulin could account for the significant decrease in the level of FBS level on the seventh day of treatment with the extract. It could be hypothesized that extract of *Bambusa vulgaris* has regenerative effect on the β-cells of pancreas in alloxan induced diabetic rats, and could be a better treatment regimen for insulin dependent diabetes mellitus.

It could be concluded that aqueous extract of *Bambusa vulgaris* has potential to reactivate damaged β-cells of pancreas, increase plasma insulin levels, and decrease the levels of blood glucose in alloxan-induced type 1 diabetes mellitus Wistar rats. Further investigations will be carried out to isolate active molecules and validate the therapeutic potential of the plant in the management of insulin dependent diabetes mellitus.

## Authors’ Contributions

MOA, JIA and FT designed the study, MOA, OSF, IOO, II, OTK, SOA and AA did the analysis. All authors prepared and approved the final manuscript.

## Acknowledgement

The authors are grateful to the Management and Staff of Ultimate Medical Diagnostic Laboratory Ibadan, Nigeria for the technical supports.

